# Innovative Methods for Prenatal Cannabis Exposure: Vapor Inhalation Chamber and Metabolite Quantification in Prairie Voles and Rats

**DOI:** 10.64898/2026.02.04.703847

**Authors:** Sophia Rogers, Casey E Hogrefe, Chun-Yi Wu, Adele M. H. Seelke, Anurupa Kar, Sabrina L. Mederos, Jessica Bond, Felisa J Carbajal, Zhenmo Yu, Melissa D. Bauman, Karen L. Bales

## Abstract

The increasing prevalence of cannabis use, including among pregnant women, highlights the critical need for a deeper understanding of prenatal cannabis exposure. This study aimed to develop a standardized cross-species inhalation exposure protocol to administer the principal psychoactive component of cannabis, Δ9-tetrahydrocannabinol (THC), to prairie voles (*Microtus ochrogaster*) and laboratory rats (*Rattus norvegicus*), and to investigate the distribution of THC in maternal and fetal tissues following prenatal exposure. Using an established e-cigarette system for delivering vaporized THC, we administered THC to pregnant prairie voles and rats. THC concentrations were measured in maternal plasma and fetal brain tissue using LC-MS/MS (Liquid Chromatography coupled with Tandem Mass Spectrometry). In both species, THC levels were compared across groups to evaluate the impact of fetal position on THC uptake. We found that THC readily crossed the placental barrier in both species, resulting in significantly higher concentrations of THC in the fetal brain within the THC-exposed groups compared to the vehicle controls. Interspecies comparison revealed higher THC concentrations in rat fetal brain tissue compared to prairie voles. No significant effects of fetal position on THC levels were found for either species. The findings confirm placental transfer of THC and reveal species-specific patterns of THC distribution. Additional studies were then carried out in voles to compare plasma and brain THC levels in maternal and virgin adult prairie voles. Maternal brain THC concentrations were significantly higher than fetal brain concentrations in prairie voles. Strong positive correlations were observed between plasma and brain THC concentrations in both maternal and virgin adult prairie voles. This study establishes a translational model for investigating prenatal cannabis exposure using an aerosolized administration method in voles compared to established methods in rats. The standardized protocol and results provide a foundation for future research into the developmental consequences of prenatal cannabis exposure and offer crucial insights for informing public health policies and clinical practices in response to the global increase in cannabis use.

## Introduction

Cannabis, a plant known for its psychoactive properties, has been used throughout history for various purposes, including medicinal, recreational, and spiritual practices (Russo, 2007; Zuardi, 2006). The main psychoactive component of cannabis is Δ9-tetrahydrocannabinol (THC), which interacts with the endocannabinoid system in the body (Mechoulam & Parker, 2013). This system consists of endogenous ligands, such as anandamide and 2-arachidonoylglycerol, which bind to G protein-coupled cannabinoid receptors (CB1 and CB2) and regulate various physiological processes (Pertwee, 2008; Lu & Mackie, 2021).

In recent years, the use of cannabis has increased significantly, becoming the third most commonly used psychoactive substance in the US after alcohol and nicotine, largely due to the legalization of its recreational and medicinal use in many states (Ross & Levy, 2023; Hall & Lynskey, 2016; Carliner et al., 2017). This increase in cannabis use has also been observed among pregnant women, with higher usage rates in the first trimester compared to the second and third trimesters, and prevalence rates ranging from 3% to 16% across various studies (Volkow et al., 2019; Young-Wolff et al., 2025a; Young-Wolff et al., 2025b; Blair et al., 2025).

The use of cannabis during pregnancy raises concerns about the potential effects of prenatal cannabis exposure on fetal development (Jansson et al., 2018). THC readily crosses the placental barrier (Bailey et al., 1987), making it important to understand its impact on the developing fetus, particularly during critical developmental periods (Schneider, 2009; Rice & Barone Jr, 2000).

A rapidly growing literature in humans gives insight on the effects of prenatal cannabis exposure (Paul et al., 2021; Sorkhou et al., 2024; Nashed et al., 2021; Young-Wolff et al., 2024; Avalos et al., 2025; Tadesse et al., 2025) including: greater risk of psychopathology, preterm delivery, low birth weight, neonatal intensive care unit (NICU) admission in newborns, physiological and neurodevelopmental consequences, gastroschisis, omphalocele, and increased anxiety risk in offspring. Human studies have explored the developmental, metabolic, and social aspects of cannabis use during pregnancy (Gunn et al., 2016; Hurd et al., 2019; Corsi et al., 2019; Jarlenski et al., 2017). Previous studies have investigated the uptake of other drugs, such as tobacco, during pregnancy (Behnke & Smith, 2013; Zhou et al., 2014; O’Connell et al., 2025). Collectively, these studies suggest that social and emotional behavior development might be particularly vulnerable to prenatal cannabis exposure, though underlying mechanisms remain poorly understood.

There is also an expanding number of animal studies showing effects of prenatal and early THC exposures (Iyer et al., 2022; Breit et al., 2022; Hussain et al., 2022; Davies et al., 2025; Roeder et al., 2024; Lei et al., 2023; Lallai et al.,2022), but animal studies examining the effects of cannabis on social behavior, particularly in prairie voles and rat models, are more scarce (Baglot et al., 2021; Frau et al., 2019; Weimar et al., 2020). The work by Baglot and colleagues is one of the few studies on prenatal THC aerosolized administration chamber exposure. Their study demonstrated that prenatal THC exposure in rats led to alterations in social behavior and neural development, highlighting the need for further investigation into the effects of prenatal cannabis exposure in highly social species.

Prairie voles (*Microtus ochrogaster*) and rats (*Rattus norvegicus*) serve as important models for studying the effects of prenatal cannabis exposure due to their well-characterized social behaviors and developmental processes (McGraw & Young, 2010; Bales & Perkeybile, 2012). Prairie voles in particular exhibit strong pair bonds and biparental care (Young & Wang, 2004), making them an attractive model for investigating the impact of prenatal drug exposure on social development. Rats have been extensively used in developmental neuroscience research (Semple et al., 2013) and provide a well-established model for studying the effects of prenatal drug exposure on brain development (Schneider, 2009). Furthermore, rats engage in rough-and-tumble play during their juvenile period, which is crucial for the development of social competence and the refinement of social, emotional, and cognitive skills (Pellis & Pellis, 2007).

The primary purpose of the current study was to further validate and build upon a standardized methodology for administering THC to rodent models through an aerosolized administration (“vaping”) chamber, evaluate maternal and fetal THC levels, and to expand this model to include a new species, prairie voles. While numerous studies have investigated prenatal cannabis exposure, there are inconsistencies in the literature regarding THC administration methods, with many relying on injections that do not accurately mimic human consumption patterns. Inhalation, particularly combustion inhalation, remains the most common route of administration for cannabis among users, due to its rapid onset of effects (Hindocha et al., 2017; Spindle et al., 2018). Our study aims to develop and validate an aerosolized administration protocol for THC administration in two rodent species: prairie voles and rats. The dosing regimen used in this study was based on previous work by Taffe et al. (2021) and Baglot et al. (2021). This approach more closely resembles human cannabis use, particularly in light of the increasing popularity of aerosolized administration among cannabis users (Budney et al., 2015). By designing and implementing this protocol, we seek to provide researchers with a more translatable method for studying the effects of prenatal cannabis exposure on fetal development.

## Methods

### Subjects

#### Prairie Voles

Prairie voles were descendants of a wild stock originally captured near Champaign, Illinois, and the colony was maintained through systematic outbreeding. The breeding, husbandry, and testing of the prairie voles were conducted at the University of California, Davis. Breeder pairs were housed in large polycarbonate cages (44×22×16cm) with aspen wood bedding (Sani-Chips) and cotton nestlets provided as nesting material. At 20 ± 1 days of age, sexually naïve male and female animals were group weaned and subsequently pair-housed in smaller (27×16×13cm) cages with either a same-sex sibling or an age-matched same-sex non-sibling. The virgin males and females used in this study were housed in (27×16×13cm) cages and were initially sexually inexperienced with no exposure to pups after weaning. Virgin males (N=6) and females (N=6) were dosed to examine the effect of sex and reproductive status on THC absorption. The voles were kept under a 14:10 light-dark cycle, with lights on at 06:00. They had ad libitum access to water and high-fiber Purina rabbit chow. The room temperature was maintained at approximately 70°F.

For the fetal exposure study, timed matings were performed to ensure exact date of conception and therefore the precise prenatal exposure day (Kenkel et al., 2019). Sexually mature males and females were placed together in a large cage with cotton for nesting material and a small amount of soiled bedding from the male’s cage to act as an olfactory stimulus. Twenty-three hours later, voles were separated from each other using a plexiglass cage divider to prevent copulation while still allowing for visual, auditory, and olfactory stimulation. The cage divider was removed two days later, allowing for mating to occur. This day was considered as the day of conception, or embryonic day 0 (E0). A total of N=4 for control pairs and a total of N=6 for exposed pairs were used for this study.

#### Rats

Sprague Dawley lab rats were obtained from Charles River Laboratories. Rats were bred, maintained, and tested at University of California, Davis. Sexually naïve male and female animals were separated upon arrival into same-sex housing pairs and allowed to acclimate to the vivarium for approximately 2 weeks. Rats were maintained on a 12:12 light-dark cycle with lights on at 07:00. Water and food were provided ad libitum. The animals were maintained and observed in large, polycarbonate cages (40×30×20cm) with corn cob bedding and paper provided for nesting material. Humidity was controlled and room temperature was maintained near 70°F. Home cages were left undisturbed beyond weekly cage changes and daily checks of food and water.

After approximately 2 weeks, breeding males were co-housed overnight with breeding females. Pregnancy was determined the next morning by the presence of a seminal plug. Upon confirmed breeding, females and males were separated and the males were singly housed. Pregnant females were paired with another female, and pregnancy was monitored by periodic measurements of weight gain. The final group size consisted of N=6 pregnant dams exposed to THC, N=2 for pregnant dams exposed to vehicle control only, and N=2 for pregnant dams as untreated controls.

### Exposure Methods

THC exposure occurred at the California National Primate Research Center (CNPRC) Respiratory Disease Center (RDC) facility. Subjects were moved to the RDC one day before the first THC exposure and were temporarily housed within the RDC during the three days of THC exposure.

Vapor Chamber Setup and Validation: Subjects were exposed to THC using the two-chamber T1 tabletop E-Vape system from La Jolla Alcohol Research, Inc. (Taffe et al., 2021; Thomas et al., 2020; Taffe et al. 2020; McLaughlin et al., 2020) (**Figure 1A**). There were two sizes of exposure chambers. Prairie vole cages fit within the smaller chamber (36.8 x 26.7 x 22.9 cm) while rat cages fit within the larger chamber (52 x 34.3 x 31.8 cm). Each chamber was validated by measuring internal pressure (rats: 18.0-18.5 cm H2O, prairie voles: 13.2-13.3 cm H2O) and flow rate to ensure consistent and accurate THC delivery to the animals. The flow rate through the chambers was measured using a mini-BUCK Primary Flow Calibrator. The flow rate for the smaller prairie vole chambers was measured at 12.34 L/min (flow rate of 6.17 L/min for each chamber). The flow rate for the bigger rat chambers was measured at 11.5 L/min (flow rate of 5.75 L/min for each chamber). Flow rates were limited using two flow meters, one regulating air input to the chamber at a flow rate of 6 standard cubic feet per hour (SCFH) of air, and one regulating the air flow from the chamber to the output pump at 12 L/min.

**Figure 1:**
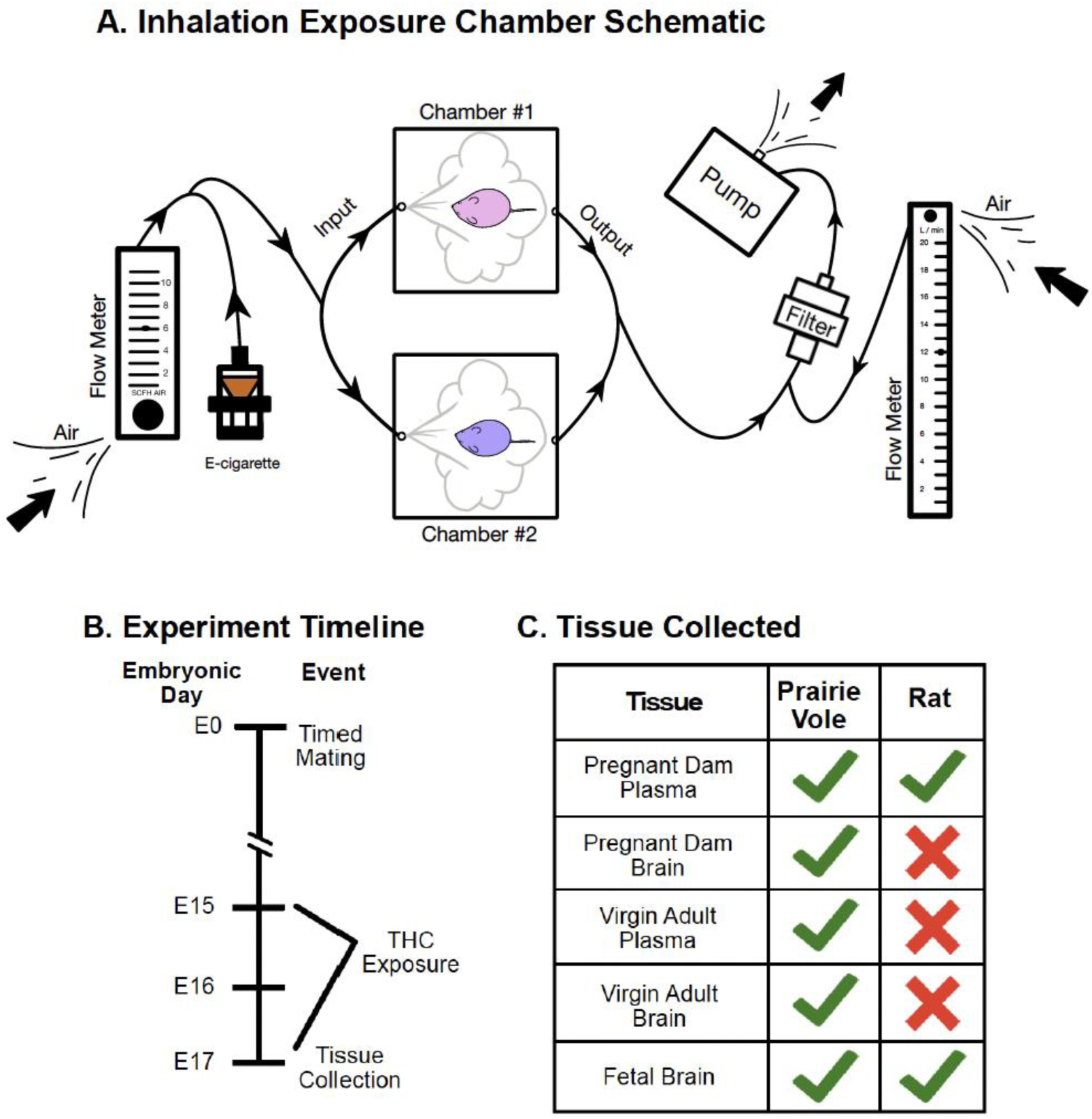
A) Schematic of apparatus. Atmospheric air is continuously pulled through the apparatus via a pump on the right side of the schematic. Air flow through the system is regulated by two flow meters, and an attached computer controls the heating of the e-cigarette and the delivery schedule of the vapor. At programmed intervals the e-cigarette releases puffs of PG or THC vapor which fills the sealed exposure chambers. Vapor output flows through a filter and is exhausted through an HVAC filtration system. B) Depicts the timeline of the experiment, including the timed mating of animals and the exposure period from embryonic day 15 (E15) to E17. Tissue collection was conducted on E17, immediately following the exposure period. C) Summarizes endpoints collected for each species.

THC e-liquid Preparation: THC was received from the National Institute on Drug Abuse (NIDA) Drug Supply Program (NDSP) as 200 mg/mL delta9-tetrahydrocannabinol (THC) in 95% ethanol solution. The THC was prepared for use in the E-Vape system by evaporating the ethanol under a stream of nitrogen gas resulting in a concentrated THC resin. The resin was dissolved in propylene glycol (PG) to a total volume twice that of the original THC-ethanol solution to achieve a concentration of 100 mg/mL. The mixture was vortexed and placed in a lukewarm water bath to ensure complete suspension. The final volume and composition of the THC-PG solution was recorded, and the mixture was wrapped in plastic wrap and foil and stored in a refrigerator at 4°C until use. The e-liquid solution was brought to room temperature prior to use and loaded into the e-cigarette (Smok® Baby Beast Brother V8 X-Baby Q2).

Vapor Chamber Experiment: The timeline of the experiment is depicted in **Figure 1B**. Both prairie voles and rats were dosed over 3 consecutive days from embryonic day 15-17 (E15-17) of the 21-day gestation. Each exposure session lasted 15 minutes with one 5-sec puff of vapor delivered every 2 minutes (Baglot, 2021). Experimental subjects were exposed to puffs of 100 mg/mL THC/PG solution, while the vehicle subjects were exposed to puffs of PG only. Untreated control rat subjects (N=2) were used as a baseline and were not placed in the apparatus or exposed to PG solutions. An outline of experimental approaches and species-specific outcomes are summarized in **Figure 1C**.

Tissue Collection: Immediately following THC exposure on Day 3, the exposed pregnant females in both species were anesthetized using isoflurane and euthanized via cervical dislocation followed by rapid decapitation. Maternal blood was collected and centrifuged to obtain plasma, which was aliquoted and stored at -20°C. Maternal vole brains were also collected and flash-frozen using dry ice. Virgin vole plasma and brain was also collected using the same procedure outline. Fetal tissues were collected and their uterine position was recorded (**Figure 2**). Fetal brains, placentas, and tissue were collected, flash-frozen using powdered dry ice, wrapped in aluminum foil, and stored in glass scintillation vials at -80°C. Prairie vole fetal tissue was used for later sex determination via PCR analysis.

**Figure 2:**
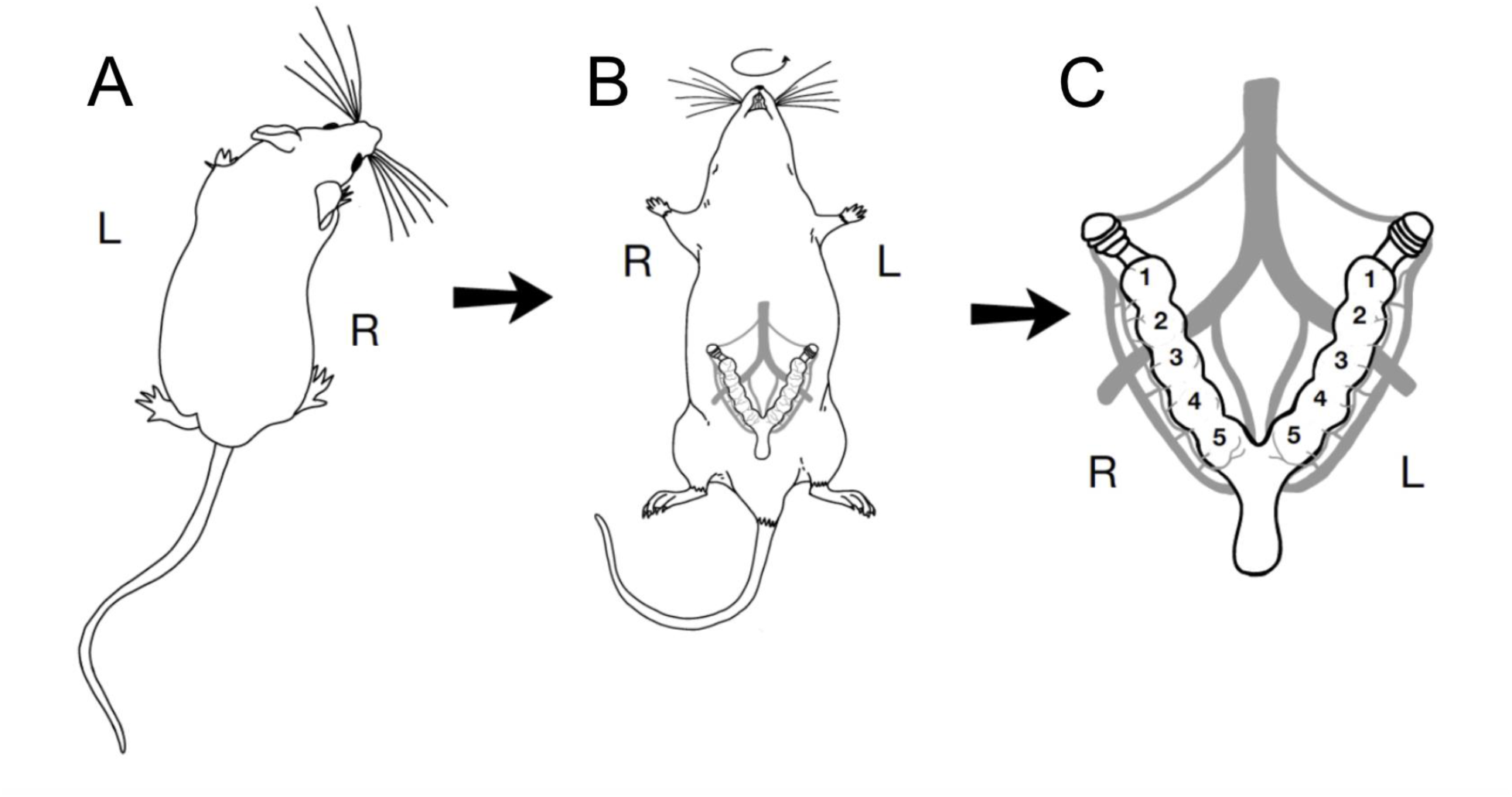
Illustration of rodent uterine anatomy. Immediately following the final session of THC exposure, dams were euthanized and fetal and maternal tissue was collected. A) Dorsal view of a behaving rodent, with left and right sides indicated. B) Rat and vole dams were placed supine to allow for collection of fetal tissue. The relative position of the uterus is shown on the ventrum, and left and right sides of the animal are maintained relative to the dorsal view. C) Detailed schematic of the anatomy of rodent uterine horns and fetal positions. The ovaries are found at the rostral end of the uterine horns, while the vaginal canal is found at the caudal point where both horns join. Fetal position is defined by placement relative to the ovaries, with “1” being the position closest to the ovary, and progressively higher numbers indicating greater distance from the ovary. Fetal subjects were given a unique identifier based on both uterine horn (left vs right) and position within the horn.

Genotyping for prairie vole fetal sex: Genomic DNA (gDNA) was extracted from fetal body tissues using a solution containing 25 mM NaOH and 0.2 mM EDTA, followed by neutralization with 40 mM Tris HCl. PCR amplification was subsequently performed using specific primers targeting the male-specific SRY gene to determine the sex of the fetus. The PCR products were analyzed by agarose gel electrophoresis to confirm the presence of the SRY gene fragment, indicative of male sex. The sequences for SRY primers for the Prairie Voles were:

forward 5′-TTATGCTGTGGTCTCGTGGTC-3′; and reverse 5′-GCA GTCTCTGTGCCTCTTGG-3′

Quantification of THC and its metabolites in plasma and brain:

Plasma and brain tissue samples were analyzed for THC and its metabolites by the UC Davis Bioanalysis and Pharmacokinetics Core Facility.

### Plasma samples

Tetrahydrocannabinol (THC) and its metabolites, 11-Hydroxy-Δ9-tetrahydrocannabinol (11-OH-THC), and 11-NOR-9-carboxy-Δ9-tetrahydrocannabinol (11-NOR-9-COOH-THC) in prairie vole and rat plasma calibrators (0.5, 1, 5, 10, 50, 100, and 500 ng/mL of each standard diluted in blank prairie vole and rat plasma with K2DTA), Quality control (QC) samples (1, 10, and 100 ng/mL of each standard diluted in blank prairie vole and rat plasma), and study samples were extracted by mixing 20 µL of the plasma with 80 µL of the internal standard solution that contains 100 ng/mL of each D_3_-THC, D_3_-11-OH-THC, and D_9_-11-NOR-9-COOH-THC in methanol (MEOH). All standards were purchased from Sigma-Aldrich (St. Louis, MO). The resulting precipitants were removed by centrifugation at 17,000 xg at room temperature for 5 minutes. Each supernatant was transferred to 96 well plate and 5 µL was injected for LC-MS/MS analysis with Waters (Milford, MA) Acquity I-class ultra-performance liquid chromatography (UPLC) hyphened with Waters Xevo TQ-S tandem triple quadrupole mass spectrometry (MS/MS). LCMS grade Water (H_2_O) with 0.1 % formic acid (FA) was used as mobile phase A (A), and acetonitrile (ACN) with 0.1% FA was used for mobile phase B (B). All solvents and reagents were purchased from Fisher (Waltham, MA). The following UPLC linear gradient was used to resolve and elute the compounds: 0∼0.25 min 40%B; 1.25 min 42.5%B; 2.00 min 50%B; 3.75 min 95%B; 3.75∼4.25 min 95%B, 4.26 min 40%B; 4.26∼5.50 min 40%B. The flow rate was maintained at 0.4 mL/min. Mobile phase A was also used for purging, and 50/50 (v/v) ACN/H_2_O was used for needle wash. Phenomenex (Torrance, CA) Kinetex 1.7 mm C18, 100 Å,100X2.1 mm was the column used for separating the compounds. The column temperature was maintained at 30° C and the autosampler temperature was maintained at 10° C. The eluted compounds were fed to the MS/MS and were first ionized with electrospray ionization at positive mode (ESI+). Those parent ions were further fragmented into daughter ions. The following Multiple reaction monitoring (MRM) mode was applied to specifically monitor the fragmentation of each compound for the quantification: THC: 315.3 -> 193.1; 11-OH-THC: 331.3 -> 193.1; 11-NOR-9-COOH-THC: 345.2 -> 193.1; D_3_-THC: 318.3 -> 196.1; D_3_-11-OH-THC: 334.3 -> 196.2; and D_9_-11-NOR-9-COOH-THC: 354.3 -> 196.8. FDA Bioanalytical Method Validation Guidance for Industry was followed to perform QC for both rat and prairie vole plasma calibration curves. The lower limit of quantification (LLOQ) QC of each test article in the corresponding matrix passed the <20% deviation (from nominal concentration) passing criteria and the rest of the QCs at all concentration levels passed the <15% deviation passing criteria. Rat plasma calibration curve also passed QC with prairie vole plasma QC samples. Therefore, rat plasma can be the surrogate matrix for prairie vole plasma, and rat calibration curves were then used to quantify both rat and prairie vole samples.

### Brain samples

THC and its metabolites in prairie vole and rat brains were quantified using the same LC-MS/MS method described above. The brains were weighed and homogenized in SPEXSampleprep (Now Cole-Parmer, Metuchen, NJ) grinding tubes pre-filled with either 2.8 mm stainless steel beads or 3 mm zirconium beads with 2-5X volume of H_2_O, assuming the density of the brain and water are equal. The tubes were shaken at 1750 rpm in SPEXsampleprep Geno/Grinder2010 for 1 minute at room temperature 3 times with a 15-second break in between. For making the brain homogenate calibrators and QC samples, the blank brains were homogenized with H_2_O and the brain homogenates were spiked with standard THC and metabolite solutions to make brain homogenate calibrators (0.5, 1, 5, 10, 50, 100, and 500 ng/mL of each standard in blank prairie vole and rat brain homogenates), and QC samples (1, 10, and 100 ng/mL of each standard diluted in blank prairie vole and rat brain homogenates). Forty microliters of each homogenate were mixed with 160 µL of the internal standard solution in methanol as mentioned above. After centrifugation at 17,000 xg at room temperature for 5 minutes, each resulting supernatant was transferred to 96 well plate and 5 µL was injected for LC-MS/MS analysis. Each rat and prairie vole brain calibration curve passed QC at all concentration levels of QC samples in each corresponding matrix. Rat brain calibration also passed QC with prairie vole brain QC samples. Therefore, rat brain homogenate can be the surrogate matrix for prairie vole brain homogenate and rat brain calibration curves were then used to quantify both rat and prairie vole brain samples.

### Statistical Analysis

All statistical analyses were conducted using RStudio (Version 2023.12.1+402; R Core Team, 2023). Data were first assessed for normality using the Shapiro-Wilk test. When data did not meet normality assumptions, nonparametric tests were performed. Group differences were analyzed using the Wilcoxon rank-sum test. Relationships between variables were evaluated using Pearson’s product-moment correlation (*cor.test*). Linear mixed-effects models were fitted using the *lme4* package. Statistical significance was defined as p < 0.05. Figures were generated using Graphpad and Prism.

## Results

### Prairie Voles

#### Prairie Vole Pregnant Dam Plasma

The Shapiro-Wilk test revealed non-normal distribution of THC concentrations in plasma (W = 0.78519, p = 0.009576) of vehicle versus THC-exposed dam prairie voles. A Wilcoxon rank-sum test showed a significant difference between dosed dam THC concentrations and vehicle dam THC concentrations for plasma (W = 24; p = 0.01421; **Figure 3A**). The average amount of plasma THC for dosed dam prairie voles was 30.65 ng/mL. For the prairie vole dam plasma, we also measured the two major metabolites of THC, 11-OH-THC and 11-NOR-9-COOH-THC (THC-COOH); **Figure 3C**. There was a significant difference between dosed dam and vehicle dam concentrations for 11-OH-THC (W = 24, p = 0.01306) and THC-COOH (W = 22, p = 0.03073) in plasma. The concentrations of the metabolites in pregnant dosed dams were significantly lower when compared to THC plasma concentrations for both 11-OH-THC (estimated -28.865, p = 0.000327; **Figure 3B**) and THC-COOH (estimated -30.181, p = 0.000216). There was a significant difference between the 11-OH-THC concentration and the THC-COOH concentration for dam prairie vole plasma (V = 21, p = 0.03603).

**Figure 3:**
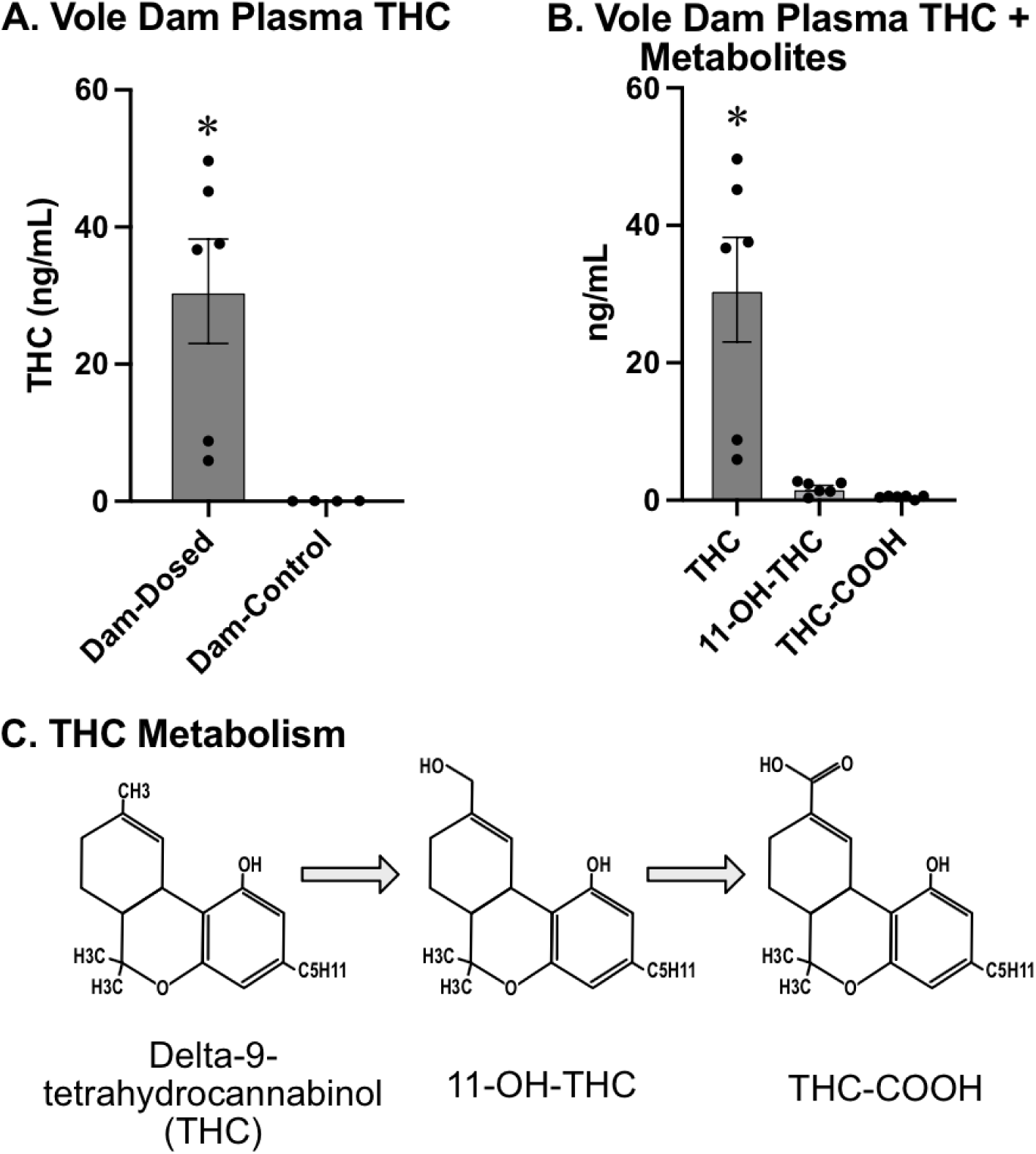
A) Concentration of THC (measured in mL of THC per ng of tissue) in the plasma of dosed prairie vole dams (left) and vehicle prairie voles dams (right). The plasma of dosed dams contained significantly higher concentrations of THC than vehicle dams. B) THC and metabolites in the dosed prairie vole dam plasma. THC-COOH concentration (right) and 11-OH-THC (center) are both significantly lower than THC (left). C) Metabolic progression of the THC molecule. Delta-9-tetrahydrocannabinol (THC) is first metabolized into 11-hydroxy-delta-9-tetrahydrocannabinol (11-OH-THC) then further metabolized into 11-Nor-9-carboxy-delta-9-tetrahydrocannabinol (THC-COOH). Ultimately, this THC-COOH is conjugated with glucuronide forming a water-soluble compound that can be easily excreted from the body. * - indicates significantly different from all other groups

#### Prairie Vole Fetal Brain

THC levels showed non-normal distribution (W = 0.75316; p = 6.455e-07) and were found to be significantly higher in dosed prairie vole fetuses compared to vehicle prairie vole fetuses (W = 399.5, p = 6.934e-08; **Figure 4A**). Across litters, the prairie vole fetal brain THC levels averaged ∼7.6 ng/g, compared with ∼30.6 ng/mL detected in the dam plasma, indicating approximately 25 % of maternal plasma concentrations.

**Figure 4:**
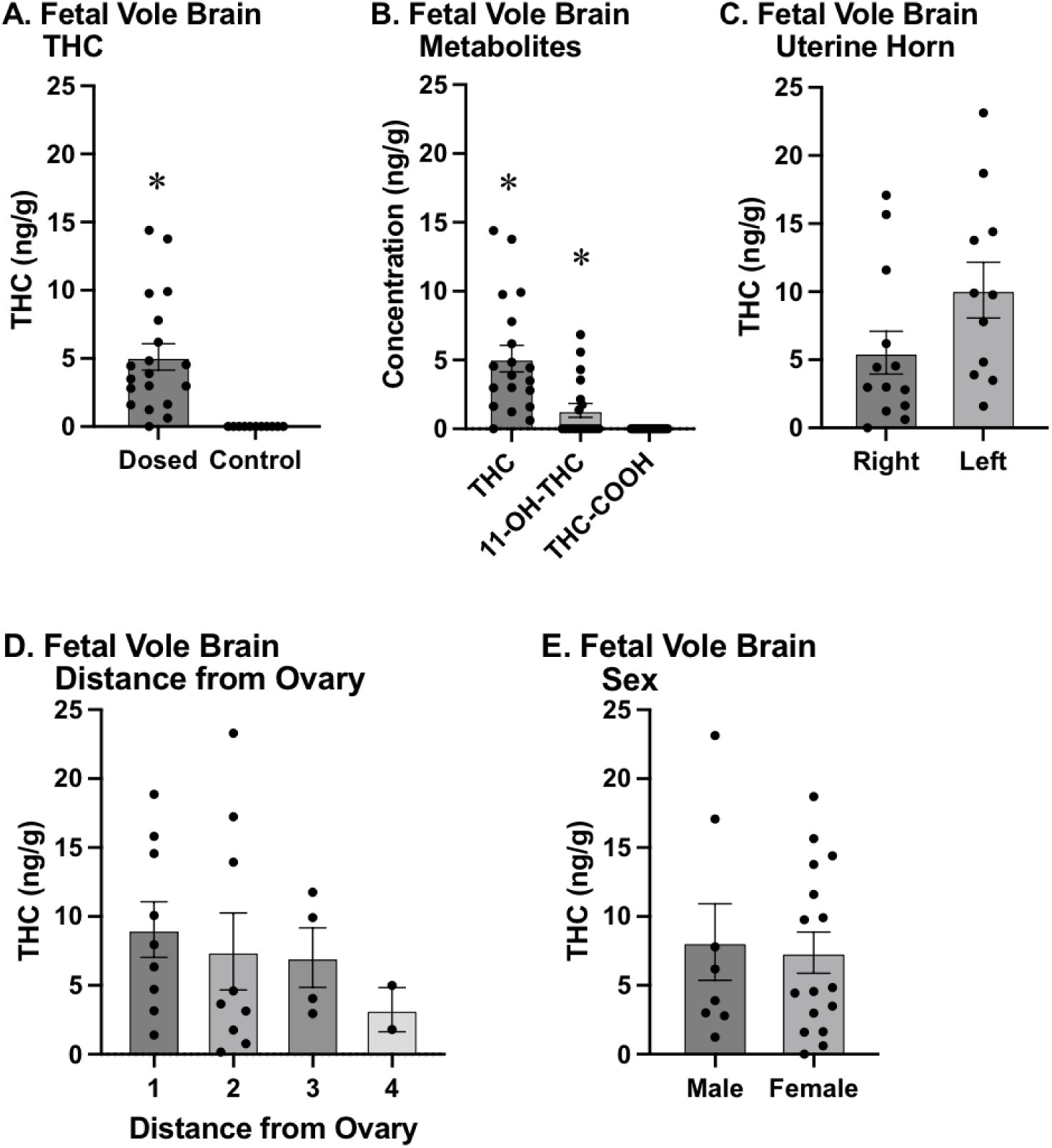
A) THC concentration (measured in ng of THC per g of tissue) in the brains of dosed (left) vs control (right) fetuses. There was significantly higher THC concentration in the brains of dosed subjects compared to control subjects, and no THC was detected in the brains of any of the control subjects. B) THC and metabolites in the fetal brain. Both THC (left) and 11-OH-THC (center) were found at significantly higher levels than COOH-THC (right) in the brains of fetuses dosed with THC. C) The concentration of THC in the brains of vole fetuses found in the left uterine horn (right) did not differ from those in the right uterine horn (left). D) Fetal position in the uterine horns was identified based on distance from the ovary, with 1 being adjacent to the ovary and higher numbers being progressively closer to the vaginal canal. THC concentration in fetal vole brains did not differ based on position within the uterine horn. E) THC concentration in the brains of dosed fetuses in relation to sexes. There was no difference in THC concentration in the brains of male and female fetal subjects. * - indicates significantly different from all other groups

In addition to THC, we measured its two major metabolites (**Figure 4B**), 11-OH-THC and THC-COOH; their presence shows that not only is THC present, but it is also being actively metabolized by the body. A Wilcoxon rank-sum test showed a significant difference in 11-OH-THC concentration between the dosed and vehicle fetal prairie vole brains (W = 306; p = 0.0008349). The THC metabolite THC-COOH was not detectable in any fetal prairie vole brains; since both groups contained only zeros, no statistical comparison was possible. A linear mixed model revealed that for dosed fetal samples, accounting for litter effects, the concentrations of 11-OH-THC and THC-COOH were significantly lower than that of THC brain concentrations, with estimated differences of -5.69 ng/g (p = 4.36e-08) and -7.63 ng/g (p = 8.34e-12). Further analysis revealed a significant difference between the concentrations of 11-OH-THC and THC-COOH in dosed fetal prairie vole brain tissue when accounting for litter effects (estimate = - 1.9413, p = 7.85e-06).

Fetal position analysis revealed no significant differences in brain THC concentrations based on uterine horn side (t = -1.377; p = 0.1844; **Figure 4C**) or specific position within the horn (t = -0.481; p = 0.6368; **Figure 4D**) after accounting for individual dam differences.

### Rats

#### Rat Pregnant Dam Plasma

THC concentrations within plasma of vehicle and untreated control dam rats were uniformly zero and no statistical comparison could be made, since there was no variability in the data. Subsequent analyses combined both vehicle and untreated control dam rat groups into a vehicle control group. A Shapiro-Wilk test revealed non-normal distribution of THC concentrations in rat dam plasma (W = 0.8381; p = 0.04187) of vehicle versus THC-exposed rats. A Wilcoxon rank-sum test showed a significant difference between dosed dam THC concentrations and vehicle control dam THC concentrations for plasma (W = 24, p = 0.01392; **Figure 5A**). The average amount of plasma THC for dosed dam rats was 88.78 ng/mL. For the rat dam plasma, we also measured 11-OH-THC and THC-COOH. For the dams, there was a significant difference between dosed dam and vehicle control dam concentrations for 11-OH-THC (W = 24, p = 0.01392) and THC-COOH (W = 24, p = 0.01142). Concentrations of the metabolites were significantly lower when compared to THC plasma concentrations for both 11-OH-THC (estimated -70.81, p = 0.000855; **Figure 5B**) and THC-COOH (estimated -74.42, p = 0.000559). There was not a significant difference between the 11-OH-THC concentration and the THC-COOH concentration for dam rat plasma (V = 19, p = 0.09349).

**Figure 5:**
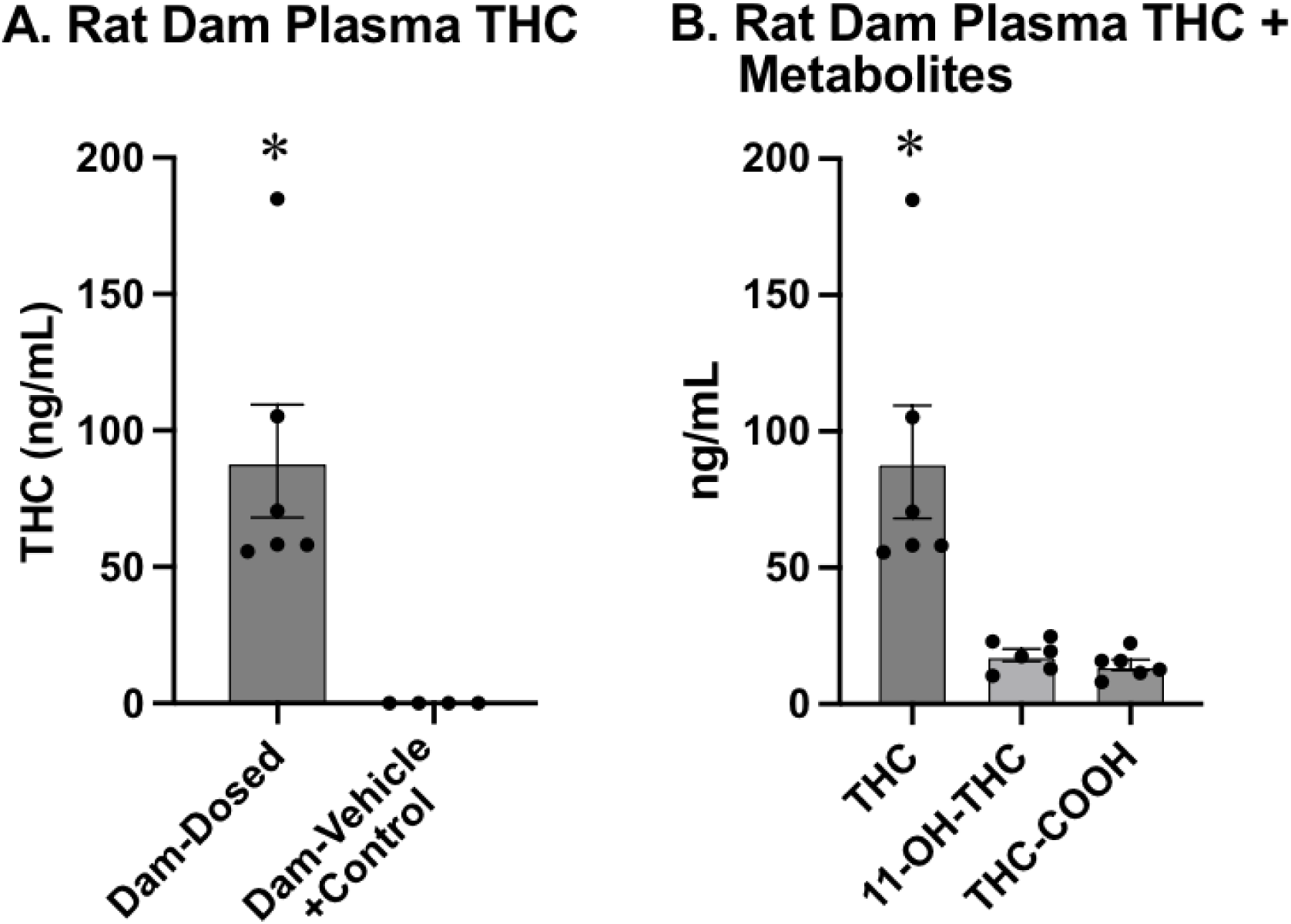
A) Concentration of THC in the plasma of dosed rat dams (left) and vehicle + control rat dams (right). The plasma of dosed rat dams contained significantly higher concentrations of THC than vehicle + control rat dams. B) THC and metabolites in the dosed rat dam plasma. THC-COOH concentration (right) and 11-OH-THC (center) are both significantly lower than THC (left). * - indicates significantly different from all other groups

#### Rat Fetal Brain

THC concentrations in fetal rat brain tissue showed non-normal distribution (W = 0.80358, p = 9.125e−13). THC concentrations for vehicle and control fetal rat brain tissue were not significantly different (p = 0.4336), and were also combined into a vehicle control group for subsequent analyses. Dosed fetal rats exhibited significantly higher brain THC concentrations compared to vehicle control fetal rats (W = 0; p = <2.2e-16; **Figure 6A**). Mean rat fetal brain THC levels were approximately 13.33 ng/g while maternal plasma levels averaged ∼ 88.78 ng/mL, indicating that fetal rat brain exposure amounted to roughly 15 % of dam plasma levels.

**Figure 6:**
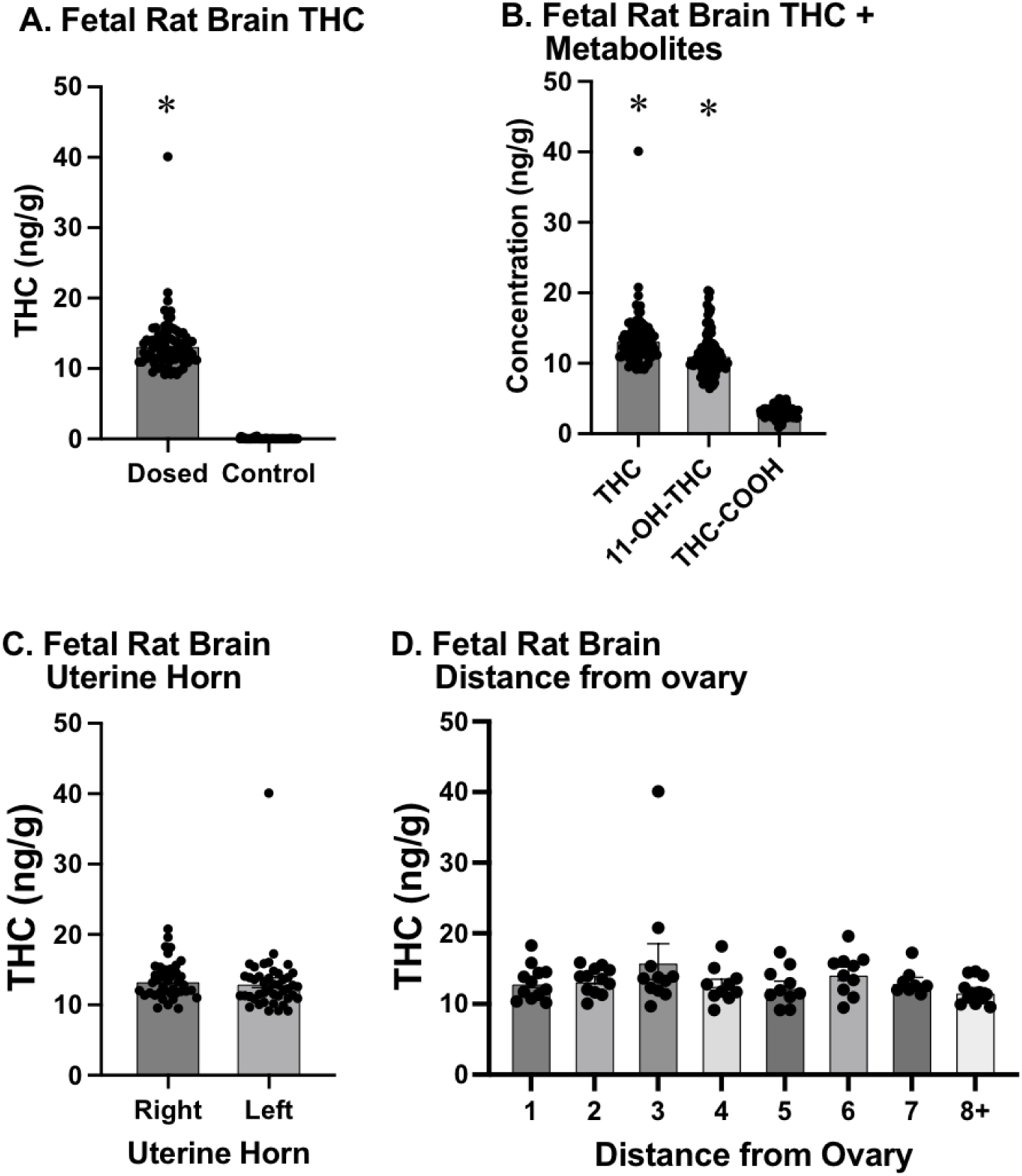
A) THC concentration (measured in ng of THC per g of tissue) in the brains of dosed (left) vs control (right) rat fetuses. There was significantly higher THC concentration in the brains of dosed subjects compared to control subjects, and no THC was detected in the brains of any of the control subjects. B) THC and metabolites in the fetal brain. Both THC (left) and 11-OH-THC (center) were found at significantly higher levels than COOH-THC (right) in the brains of fetuses dosed with THC. C) The concentration of THC in the brains of rat fetuses found in the left uterine horn (right) did not differ from those in the right uterine horn (left). D) Fetal position in the uterine horns was identified based on distance from the ovary, with 1 being adjacent to the ovary and higher numbers being progressively closer to the vaginal canal. THC concentration in fetal rat brains did not differ based on position within the uterine horn. * - indicates significantly different from all other groups

In addition to THC, we measured its two major metabolites in fetal rats. A Wilcoxon rank-sum test showed a significant difference in 11-OH-THC concentration between the dosed and vehicle fetal rat brains (W = 0; p = <2.2e-16). A Wilcoxon rank-sum test showed a significant difference in THC-COOH concentration between the dosed and vehicle fetal rat brains (W = 3; p = <2.2e-16). A comparison using linear mixed models was conducted to examine the concentrations of THC and its metabolites, 11-OH-THC and THC-COOH, for dosed rat fetal samples and indicated that the concentrations of 11-OH-THC (estimate = -2.1229, p = 2.68e-07; **Figure 6B**) and THC-COOH (estimate = -10.2653, p = <2e-16) were significantly lower than that of THC brain concentrations, when accounting for litter effects. The concentrations of 11-OH-THC and THC-COOH in fetal rat brain tissue were significantly different from each other with THC-COOH lower than 11-OH-THC (estimate = 8.1424, p = <2e-16).

No significant difference was found between left and right uterine horn sides (p = 0.949; **Figure 6C**) or between specific positions within the horn (p = 0.579; **Figure 6D**) accounting for individual dam variability.

#### Interspecies Comparison

A Wilcoxon rank-sum test revealed significantly higher plasma THC concentrations in rat dams compared to prairie vole dams (W = 36; p = 0.005075; **Figure 7A**). For the metabolites of THC in plasma, the 11-OH-THC concentration between dosed rat dams and dosed prairie vole dams was significant (W = 36; p = 0.005075). The THC-COOH concentration for the dams of both species was also found to be significantly different (W = 36; p = 0.005075).

**Figure 7:**
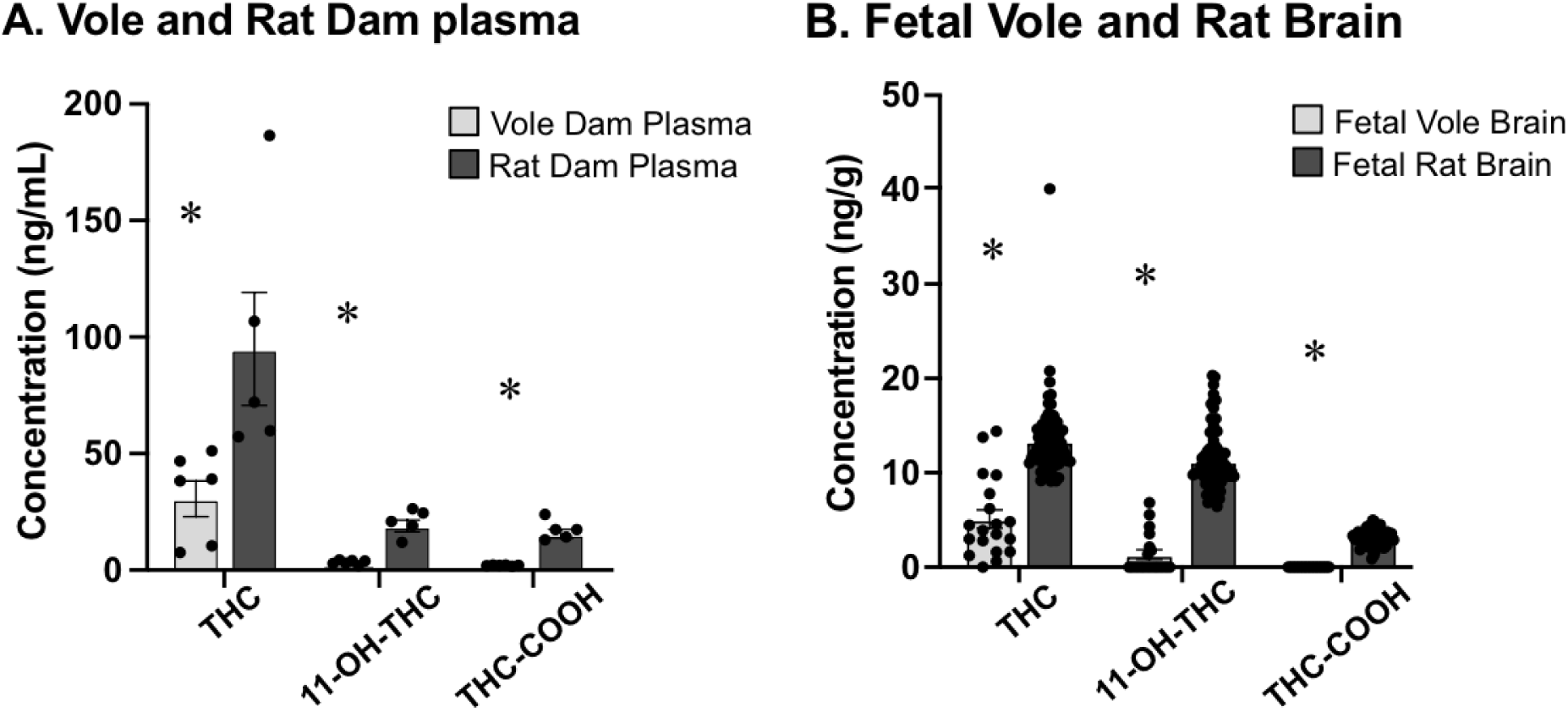
Direct comparison of THC concentration in adult (A) and fetal (B) rats and voles. A) Plasma THC, 11-OH-THC and THC-COOH concentration were all significantly higher in rat dams (dark grey) than vole dams (light grey). B) Brain THC, 11-OH-THC and THC-COOH concentration were all significantly higher in rat fetal brains (dark grey) than vole fetal brains (light grey) * - indicates significant difference between species

Fetal brain THC concentrations were found to be significantly higher in THC-exposed fetal rat brain tissue compared to THC-exposed fetal voles (W = 1569; p = 6.045e-05; **Figure 7B**). After running a Wilcoxon rank-sum test for the dosed fetal brains of the rats and prairie voles, the metabolites also showed a significant difference between species in concentration for the 11-OH-THC (W = 2039; p = 9.124e-14) and the THC-COOH (W = 2040, p = 6.601e-14).

### Additional Prairie Vole Results

#### Prairie Vole Pregnant Dam Brain vs. Fetal Brain

The Shapiro-Wilk test revealed a normal distribution of THC concentrations in dam prairie vole brains between vehicle and dosed dams (W = 0.87898, p = 0.127). A Wilcoxon rank-sum test showed a significant difference between dosed dam THC concentrations and vehicle dam THC concentrations for the brain (W = 24; p = 0.01142; **Figure 8A**). Maternal brain THC concentrations were found to be significantly higher than fetal THC levels in a linear mixed-effects model accounting for individual dam variability (estimate = 24.6, p = 1.05e-09; **Figure 8B**). There was also a significant difference in both the metabolites of THC in relation to dosed dam brain and dosed fetal brain, 11-OH-THC (estimate 8.775, p = 6.53e-08) and THC-COOH (estimate 0.106198, p = 0.0439).

**Figure 8:**
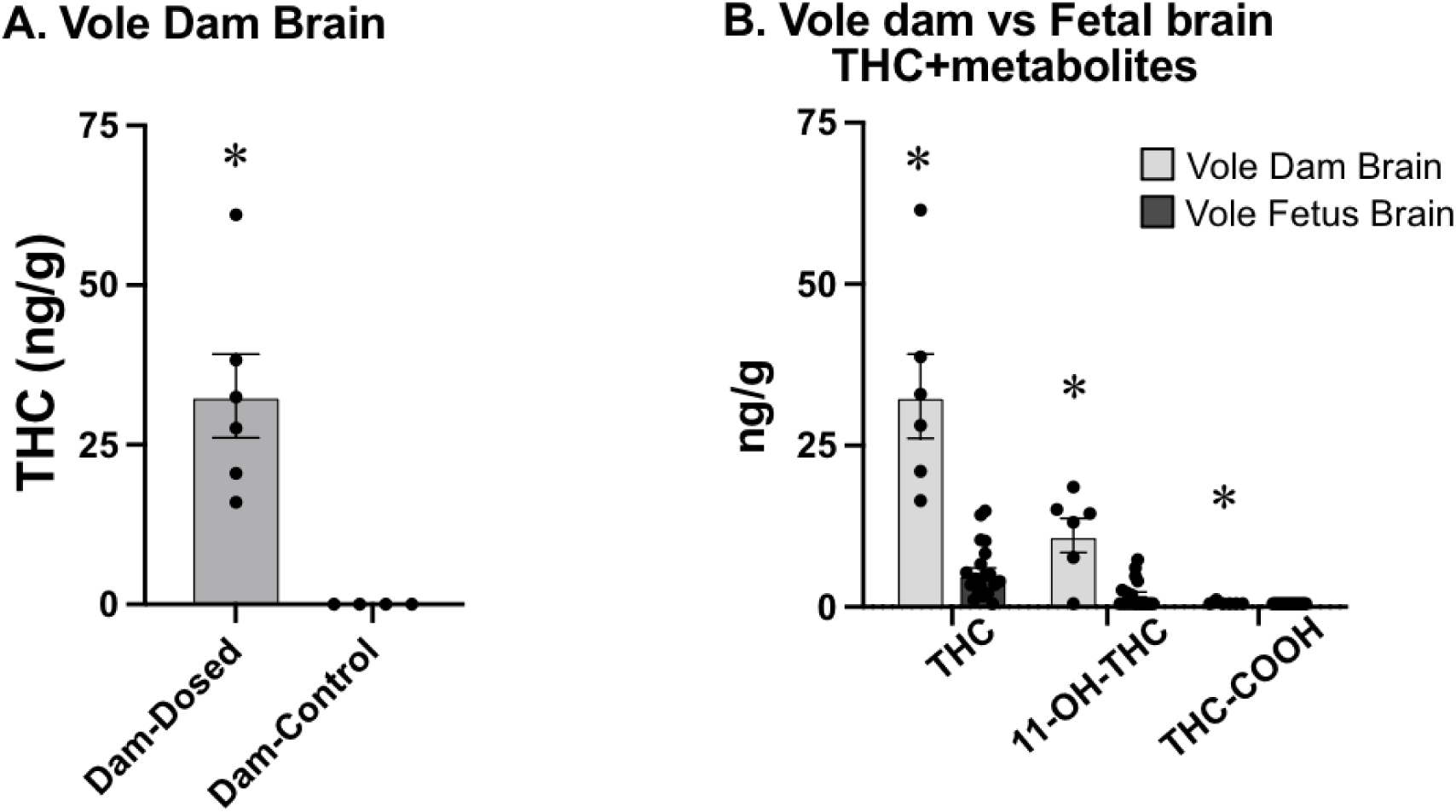
A) Concentration of THC in the brains of dosed dams (left) and vehicles (right). The dosed dam brains were significantly higher than the vehicle dam brains. B) Concentration of THC (left), 11-OH-THC (middle), and THC-COOH (right) in the brains of prairie vole dams and fetal prairie voles. The concentration for the dams was significantly higher than fetuses for THC and both its metabolites. * - indicates significantly different from all other groups

#### Prairie Vole Adult Brain and Plasma

A subsequent Wilcoxon rank-sum test did not reveal a sex difference between male and female virgin prairie voles for brain THC concentration (W = 13; p = 0.1939) or for plasma THC concentration (W = 7; p = 0.8852). Since there were no sex differences, only female virgins were used for subsequent analyses because they more closely approximate the experimental subjects. No significant difference in brain THC concentrations was found between dosed dams and female virgins (W = 15; p = 0.594; **Figure 9A**). For the adult dam brains and virgin female brains, we also measured the two major metabolites of THC, 11-OH-THC and THC-COOH, and found some significant differences between THC and its metabolites (**Figure 9B**). There was a significant difference between the dosed and vehicle dams for 11-OH-THC (W = 22, p = 0.03073), but not a significant difference between dosed and vehicle dams for THC-COOH (W = 14, p = 0.5403). For the dosed dams, the concentrations of the metabolites were significantly lower when compared to THC brain concentrations for both 11-OH-THC (estimated -21.56 ng/g, p = 0.00197) and THC-COOH (estimated -32.535 ng/g, p = 4.67e-05). For the dosed female virgins, the metabolite concentrations were also significantly lower than THC for both 11-OH-THC (estimated -19.925 ng/g, p = 0.000553) and THC-COOH (estimated -25.511 ng/g, p = 9.10e-05) compared to THC brain concentrations. Concentrations of THC and its metabolites did not differ significantly between dams and virgin females. However, plasma THC concentrations differed significantly between female virgin and dam prairie voles (W = 1; p = 0.02518; **Figure 9C**).

**Figure 9:**
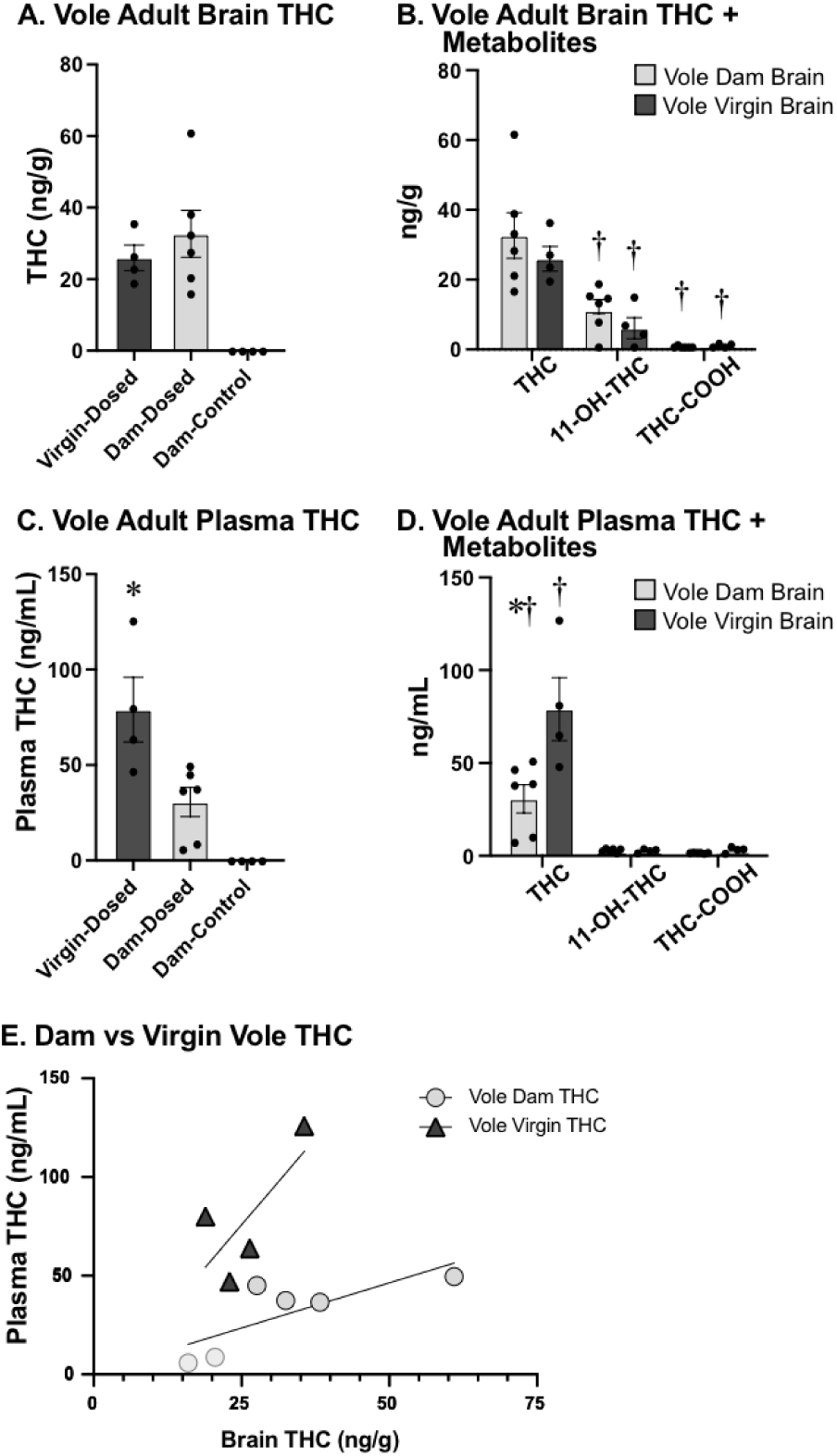
A) Concentration of THC in the brains of dosed virgin female voles (left), dosed dams (center), and vehicle dams (right). The brains of the dosed dams contained significantly higher concentrations of THC than those of control dams. B) Concentration of THC (left), 11-OH-THC (middle), and THC-COOH (right) in the adult brains of prairie vole dams and female virgins. There was a significant difference in THC and metabolite concentrations within a group, but no significant difference between dam and virgin female dosed brains. C) The concentration of THC in the plasma of dosed virgin female voles (left), dosed dams (center), and control dams (right). The plasma of dosed dams contained significantly lower concentrations of THC than virgin females, and significantly higher concentrations of THC than control dams. D) THC and metabolite concentrations in the plasma of dosed vole dams. 11-OH-THC concentration (center) and THC-COOH concentration (right) were both significantly lower than THC (left) in the plasma of dams. There was not a significant difference between 11-OH-THC and THC-COOH. E) Correlation between brain (x-axis; ng/g) and plasma (y-axis; ng/mL) THC concentrations in dams (clear circles) and virgin females (black triangles). * - indicates a significant difference between dams and virgin females. + = difference in THC and metabolite concentrations within a group.

#### Prairie Vole Virgin Plasma Metabolites

Additionally, for the virgin female plasma, we measured 11-OH-THC and THC-COOH. For the dosed female virgins, the metabolite concentrations for plasma were also significantly lower than THC for both 11-OH-THC (estimated -77.415 ng/g, p = 0.000339) and THC-COOH (estimated -76.940 ng/g, p = 0.000354; **Figure 9D**). The metabolite concentrations, 11-OH-THC and THC-COOH, were not significantly different from each other for prairie vole virgin female plasma (V = 4, p = 0.8551). Concentrations of the metabolites of THC did not differ significantly between dams and virgin females, 11-OH-THC (W = 13, p = 0.9151) and THC-COOH (W = 5.5, p = 0.1995).

#### Prairie Voles Adult Brain vs Plasma

A positive correlation, although not significant, was observed between plasma and brain tissue THC concentrations for the dams (r = 0.785, p = 0.065; **Figure 9E**) and female virgins (r = 0.737, p = 0.2635).

#### Prairie Vole Fetal Sex

For the fetal dosed prairie voles, extracted fetal body tissues were used to determine the sex of the fetus. There was no significant difference in the THC concentration of fetal brains between male and female fetuses after accounting for litter effects (estimate = 1.058, p = 0.497; **Figure 4E**). There were also no significant differences between male and female fetal brains for 11-OH-THC after accounting for litter effects (estimate = 0.2786, p = 0.6179). THC-COOH was undetectable in all fetal prairie vole samples, so no statistical comparison could be run.

## Discussion

These studies utilized established e-cigarette vapor technology protocols (Taffe et al., 2021, Nguyen et al., 2016, Nguyen et al., 2018, Javadi-Paydar et al., 2019) to deliver THC to pregnant vole and rat dams. While previous studies have utilized this translationally relevant delivery in rats (Ginder et al., 2024, Freels et al., 2020, Preteroti et al., 2023, Baglot et al., 2021), we are unaware of comparable studies in voles.

In this series of experiments, we demonstrated the efficacy of prenatal dosing of rodents with inhaled THC. We established that THC exposure within a vapor chamber will enter into the maternal blood in both species, and that THC will also cross the placental barrier into fetuses.

Furthermore, the presence of THC metabolites in fetal brain tissue demonstrates that the THC is actively metabolized by the fetus. Finally, we demonstrated that there are species-specific differences in the concentration of THC in both the dams and offspring of rats and prairie voles. Together, these results establish the vapor chamber as a reliable method of prenatal THC exposure and demonstrate the importance of carefully choosing the appropriate model species for a research question.

Prior studies have shown that the primary psychoactive compound in marijuana, Delta-9-tetrahydrocannabinol (THC), crosses the placental barrier quickly, whereas its primary metabolite (THC-COOH) does not (Bailey et al., 1987). Our data supports this idea, as the concentration of THC-COOH was significantly lower than THC for both the fetal prairie voles and fetal rats. The placenta appears to reduce fetal exposure to marijuana, as evidenced by research in various animal models, which has documented that fetal THC concentrations are consistently lower than maternal concentrations (Behnke & Smith, 2013). Our findings in prairie voles corroborate these earlier studies, demonstrating significantly higher THC concentrations in maternal brain tissue compared to fetal brain tissue. This difference suggests that while THC does cross the placental barrier, there may be additional protective mechanisms in place that limit fetal exposure (Kumar et al., 2023). However, it is crucial to note that despite this reduction, THC levels in dosed prairie vole fetuses were still significantly higher than in vehicle controls, indicating that prenatal exposure does occur and could potentially impact fetal development. Another important finding was that, while there were no significant differences in brain THC concentrations between prairie vole dosed dams and female virgins, there was a significant difference between the plasma THC concentrations of female virgin and dam prairie voles. A possible explanation could be that the fetuses are absorbing some of the THC circulating in the blood of the dam prairie voles. A study by the Lo group (Shorey-Kendrick et al., 2023), reported placental dysfunction and altered DNA methylation following prenatal THC exposure in rhesus monkeys, indicating the potential of additional effects of THC on the placenta directly. In rats, our results similarly showed significantly higher THC concentrations in dosed fetal brain tissue compared to vehicle controls. The absence of differences between left and right uterine horn locations and uterine position in both rats and voles supports the finding that fetal position does not significantly impact THC exposure. This finding is important for understanding the uniformity of THC distribution across multiple fetuses and may have implications for studying the effects of prenatal THC exposure on litter-wide outcomes.

Our exposure paradigm is based on Baglot et al. (2021), which reported that repeated THC vapor inhalation in pregnant Sprague Dawley rats resulted in higher THC concentrations in the maternal blood compared to the fetal brain, with fetal brain THC concentrations at about one third of dam blood levels. The authors concluded that the inhalation route produced lower fetal brain exposure than the more commonly used injection protocols. Similar to the Baglot results, prairie vole fetal brain THC concentrations reached only a fraction of the prairie vole dam circulating levels, averaging ∼25 % of maternal plasma concentrations. For the rats, the mean fetal brain THC concentrations corresponded to ∼ 15 % of maternal plasma concentrations, also reflecting restricted dam-fetal THC transfer, but at a higher level. The difference in the Baglot et al. report of ∼30% rat fetal brain THC relative to maternal rat blood and our ∼15% rat THC ratio could be due to a variety of slight variations in procedures. It is important to note that Baglot et al. exposed dams to THC for 12 consecutive days and collected samples 15 minutes after the final dose, whereas our dams received 3 days of exposure and samples were collected immediately after the final session, which may contribute to the lower ∼15% ratio observed in our rat cohort. Certain methodological details, including whether whole blood or plasma was analyzed or whether the fetal brains were assayed as a whole-brain tissue or a specific region could also have contributed to the differences. Potential variation in the way the THC was quantified may further explain the fetal-dam ratios observed across studies. Our observed average rat dam plasma concentration (∼88 ng/mL) was also higher than Baglot et al. (∼65 ng/mL) and other reported in studies focusing on motor effects (∼20 ng/mL; Hussain et al., 2022, Breit et al., 2020; Breit et al., 2022), which may be attributable to differences in sampling time as those studies also collected 15-20 min post-exposure.

Our use of an aerosolized administration chamber for THC administration represents a unique methodological process. This approach more closely mimics human consumption patterns, particularly given the increasing popularity of vaping among cannabis users. By standardizing this method across two rodent species, we provide a more translational model for studying the effects of prenatal cannabis exposure. While our study provides valuable insights into the distribution of THC in maternal and fetal tissues, it also highlights the need for further research. Future studies should investigate the long-term developmental consequences of the observed THC exposure, particularly focusing on neurobehavioral outcomes (Halbout et al., 2023). Additionally, exploring the mechanisms behind the species differences in THC concentrations could provide valuable insights into the factors influencing prenatal THC exposure. The interspecies comparison revealing higher THC concentrations for rats compared to prairie voles for both fetal brain tissue and dam plasma is particularly intriguing. This difference could be attributed to various factors, including species-specific placental structure, metabolism, or distribution of THC. Furthermore, even animals that are closely related, like those in family *Muridae* of class *Rodentia,* show different patterns of development, anatomy, and behavior (Krubitzer et al., 2011). In this experiment, we chose to examine rats and prairie voles due to their specific behaviors that are seen in humans, but not commonly expressed in other laboratory rodents: juvenile play and pair bond formation. Our ultimate goal is to understand how prenatal exposure to THC impacts the development of complex social behaviors later in life, and the well-established social behaviors exhibited by rats and prairie voles provide an excellent experimental model that will help us extrapolate findings to human scenarios. Both species had plasma THC concentrations that are within the range of human THC plasma concentrations, with plasma THC reaching ∼18-110 ng/mL in humans (Schwope et al., 2011).

In conclusion, our study shows that prenatal THC exposure via aerosolized administration leads to significant concentrations of THC in the fetal brain of both prairie voles and rats, although these levels are lower compared to those found in dam tissue. The standardized aerosolized administration protocol used in this study provides a robust method for future investigations into the effects of prenatal cannabis exposure. If the use of cannabis continues to increase as it has in past years, including among pregnant women, understanding the implications of prenatal exposure becomes increasingly critical (Young-Wolff et al., 2024; Mattingly et al., 2024).

## Acknowledgements

P51 OD011107 to Simon Atkinson

UC-Davis Cannabis and Hemp Research Center

UC-Davis Academic Senate

MDB received a pilot grant from the UC-Davis Academic Senate

KLB received a pilot grant from the UC-Davis Cannabis and Hemp Research Center

## Notes

### Competing Interest Statement

The authors have declared no competing interest.

## References

Avalos, L. A., Adams, S. R., Alexeeff, S. E., Oberman, N. R., Does, M. B., Steuerle, K. R., Ansley, D. R., Castellanos, C. L., Padon, A. A., Silver, L. D., & Young-Wolff, K. C. (2025). Maternal Prenatal Cannabis Use and Major Structural Birth Defects. Birth Defects Research, 117(6), e2492. 10.1002/bdr2.2492

Baglot, S. L., VanRyzin, J. W., Marquardt, A. E., Aukema, R. J., Petrie, G. N., Hume, C., Reinl, E. L., Bieber, J. B., McLaughlin, R. J., McCarthy, M. M., & Hill, M. N. (2022). Maternal-fetal transmission of delta-9-tetrahydrocannabinol (THC) and its metabolites following inhalation and injection exposure during pregnancy in rats. Journal of Neuroscience Research, 100(3), 713–730. 10.1002/jnr.24992

Bailey, J. R., Cunny, H. C., Paule, M. G., & Slikker, W. (1987). Fetal disposition of Δ9-tetrahydrocannabinol (THC) during late pregnancy in the rhesus monkey. Toxicology and Applied Pharmacology, 90(2), 315–321. 10.1016/0041-008X(87)90338-3

Bales, K. L., & Perkeybile, A. M. (2012). Developmental experiences and the oxytocin receptor system. Hormones and Behavior, 61(3), 313–319. 10.1016/j.yhbeh.2011.12.013

Behnke, M., & Smith, V. C. (2013). Prenatal Substance Abuse: Short- and Long-term Effects on the Exposed Fetus. Pediatrics, 131(3), e1009–e1024. 10.1542/peds.2012-3931

Blair, L. M., Shukla, M., Kurzer, J. A. M. J., Schilt-Solberg, M., Strickland, B. A., Akter, S., Fend, D., Hamann, K., & Ashford, K. (2025). Cannabis use patterns, motivations, and reasons for abstinence in pregnancy. Frontiers in Psychiatry, 16, 1613324. 10.3389/fpsyt.2025.1613324

Breit, K. R., Rodriguez, C. G., Lei, A., Hussain, S., & Thomas, J. D. (2022). Effects of prenatal alcohol and delta-9-tetrahydrocannabinol exposure via electronic cigarettes on motor development. *Alcoholism*, Clinical and Experimental Research, 46(8), 1408–1422. 10.1111/acer.14892

Breit, K. R., Rodriguez, C. G., Lei, A., & Thomas, J. D. (2020). Combined vapor exposure to THC and alcohol in pregnant rats: Maternal outcomes and pharmacokinetic effects. Neurotoxicology and Teratology, 82, 106930. 10.1016/j.ntt.2020.106930

Budney, A. J., Sargent, J. D., & Lee, D. C. (2015). Vaping cannabis (marijuana): Parallel concerns to e-cigs? Addiction, 110(11), 1699–1704. 10.1111/add.13036

Calapai, F., Cardia, L., Sorbara, E. E., Navarra, M., Gangemi, S., Calapai, G., & Mannucci, C. (2020). Cannabinoids, Blood–Brain Barrier, and Brain Disposition. Pharmaceutics, 12(3), 265. 10.3390/pharmaceutics12030265

Carliner, H., Brown, Q. L., Sarvet, A. L., & Hasin, D. S. (2017). Cannabis use, attitudes, and legal status in the U.S.: A review. Preventive Medicine, 104, 13–23. 10.1016/j.ypmed.2017.07.008

Corsi, D. J., Walsh, L., Weiss, D., Hsu, H., El-Chaar, D., Hawken, S., Fell, D. B., & Walker, M. (2019). Association Between Self-reported Prenatal Cannabis Use and Maternal, Perinatal, and Neonatal Outcomes. JAMA, 322(2), 145–152. 10.1001/jama.2019.8734

Davies, S., Dominguez, Z. M. & Maxwell, J. R. (2025). Preclinical Model of Prenatal Delta-9-Tetrahydrocannabinol Exposure to Assess Its Impact on Neurodevelopmental Outcomes. Journal of Visualized Experiments (JoVE), 216, e68119. 10.3791/68119

Frau, R., Miczán, V., Traccis, F., Aroni, S., Pongor, C. I., Saba, P., Serra, V., Sagheddu, C., Fanni, S., Congiu, M., Devoto, P., Cheer, J. F., Katona, I., & Melis, M. (2019). Prenatal THC exposure produces a hyperdopaminergic phenotype rescued by pregnenolone. Nature Neuroscience, 22(12), 1975–1985. 10.1038/s41593-019-0512-2

Freels, T. G., Baxter-Potter, L. N., Lugo, J. M., Glodosky, N. C., Wright, H. R., Baglot, S. L., Petrie, G. N., Yu, Z., Clowers, B. H., Cuttler, C., Fuchs, R. A., Hill, M. N., & McLaughlin, R. J. (2020a). Vaporized Cannabis Extracts Have Reinforcing Properties and Support Conditioned Drug-Seeking Behavior in Rats. Journal of Neuroscience, 40(9), 1897–1908. 10.1523/JNEUROSCI.2416-19.2020

Freels, T. G., Baxter-Potter, L. N., Lugo, J. M., Glodosky, N. C., Wright, H. R., Baglot, S. L., Petrie, G. N., Yu, Z., Clowers, B. H., Cuttler, C., Fuchs, R. A., Hill, M. N., & McLaughlin, R. J. (2020b). Vaporized Cannabis Extracts Have Reinforcing Properties and Support Conditioned Drug-Seeking Behavior in Rats. The Journal of Neuroscience: The Official Journal of the Society for Neuroscience, 40(9), 1897–1908. 10.1523/JNEUROSCI.2416-19.2020

Ginder, D. E., Weimar, H. V., Lindberg, J. E. M., Fisher, Z. D. G., Lim, M. M., Peters, J. H., & McLaughlin, R. J. (2024). Maternal cannabis use alters excitatory inputs to corticostriatal efferent neurons in rat offspring (p. 2024.03.03.583210). bioRxiv. 10.1101/2024.03.03.583210

Gunn, J. K. L., Rosales, C. B., Center, K. E., Nuñez, A., Gibson, S. J., Christ, C., & Ehiri, J. E. (2016). Prenatal exposure to cannabis and maternal and child health outcomes: A systematic review and meta-analysis. BMJ Open, 6(4), e009986. 10.1136/bmjopen-2015-009986

Gutierrez, A., Creehan, K. M., Turner, M. L., Tran, R. N., Kerr, T. M., Nguyen, J. D., & Taffe, M. A. (2021). Vapor exposure to Δ9-tetrahydrocannabinol (THC) slows locomotion of the Maine lobster (Homarus americanus). Pharmacology, Biochemistry, and Behavior, 207, 173222. 10.1016/j.pbb.2021.173222

Halbout, B., Hutson, C., Hua, L., Inshishian, V., Mahler, S. V., & Ostlund, S. B. (2023). Long-term effects of THC exposure on reward learning and motivated behavior in adolescent and adult male rats. Psychopharmacology, 240(5), 1151–1167. 10.1007/s00213-023-06352-4

Hall, W., & Lynskey, M. (2016). Evaluating the public health impacts of legalizing recreational cannabis use in the United States. Addiction, 111(10), 1764–1773. 10.1111/add.13428

Hindocha, C., Freeman, T. P., & Curran, H. V. (2017). Anatomy of a Joint: Comparing Self-Reported and Actual Dose of Cannabis and Tobacco in a Joint, and How These Are Influenced by Controlled Acute Administration. Cannabis and Cannabinoid Research, 2(1), 217–223. 10.1089/can.2017.0024

Huestis, M. A. (2007). Human Cannabinoid Pharmacokinetics. Chemistry & Biodiversity, 4(8), 1770–1804. 10.1002/cbdv.200790152

Hurd, Y. L., Manzoni, O. J., Pletnikov, M. V., Lee, F. S., Bhattacharyya, S., & Melis, M. (2019). Cannabis and the Developing Brain: Insights into Its Long-Lasting Effects. Journal of Neuroscience, 39(42), 8250–8258. 10.1523/JNEUROSCI.1165-19.2019

Hussain, S., Breit, K. R., & Thomas, J. D. (2022). The effects of prenatal nicotine and THC E-cigarette exposure on motor development in rats. Psychopharmacology, 239(5), 1579–1591. 10.1007/s00213-022-06095-8

Iyer, P., Niknam, Y., Campbell, M., Moran, F., Kaufman, F., Kim, A., Sandy, M., & Zeise, L. (2022). Animal evidence considered in determination of cannabis smoke and Δ9 - tetrahydrocannabinol (Δ9 -THC) as causing reproductive toxicity (developmental endpoint); Part II. Neurodevelopmental effects. Birth Defects Research, 114(18), 1155–1168. 10.1002/bdr2.2084

Jansson, L. M., Jordan, C. J., & Velez, M. L. (2018). Perinatal Marijuana Use and the Developing Child. JAMA, 320(6), 545–546. 10.1001/jama.2018.8401

Jarlenski, M., Koma, J. W., Zank, J., Bodnar, L. M., Bogen, D. L., & Chang, J. C. (2017). Trends in perception of risk of regular marijuana use among US pregnant and nonpregnant reproductive-aged women. American Journal of Obstetrics & Gynecology, 217(6), 705–707. 10.1016/j.ajog.2017.08.015

Javadi-Paydar, M., Kerr, T. M., Harvey, E. L., Cole, M., & Taffe, M. A. (2019). Effects of nicotine and THC vapor inhalation administered by an electronic nicotine delivery system (ENDS) in male rats. Drug and Alcohol Dependence, 198, 54–62. 10.1016/j.drugalcdep.2019.01.027

Kenkel, W. M., Perkeybile, A.-M., Yee, J. R., Pournajafi-Nazarloo, H., Lillard, T. S., Ferguson, E. F., Wroblewski, K. L., Ferris, C. F., Carter, C. S., & Connelly, J. J. (2019). Behavioral and epigenetic consequences of oxytocin treatment at birth. Science Advances, 5(5), eaav2244. 10.1126/sciadv.aav2244

Krubitzer, L., Campi, K. L., & Cooke, D. F. (2011). All Rodents Are Not the Same: A Modern Synthesis of Cortical Organization. Brain, Behavior and Evolution, 78(1), 51. 10.1159/000327320

Kumar, A. R., Sheikh, E. D., Monson, J. W., Ligon, S. E., Talley, R. L., Dornisch, E. M., Howitz, K. J., Damicis, J. R., Ieronimakis, N., & Unadkat, J. D. (2023). Understanding the Mechanism and Extent of Transplacental Transfer of (-)-Δ9 -Tetrahydrocannabinol (THC) in the Perfused Human Placenta to Predict In Vivo Fetal THC Exposure. Clinical Pharmacology and Therapeutics, 114(2), 446–458. 10.1002/cpt.2964

Lallai, V., Manca, L., Sherafat, Y., & Fowler, C. D. (2022). Effects of Prenatal Nicotine, THC, or Co-Exposure on Cognitive Behaviors in Adolescent Male and Female Rats. Nicotine & Tobacco Research: Official Journal of the Society for Research on Nicotine and Tobacco, 24(8), 1150–1160. 10.1093/ntr/ntac018

Lei, A., Breit, K. R., & Thomas, J. D. (2023). Prenatal alcohol and tetrahydrocannabinol exposure: Effects on spatial and working memory. Frontiers in Neuroscience, 17, 1192786. 10.3389/fnins.2023.1192786

Lu, H.-C., & Mackie, K. (2021). Review of the Endocannabinoid System. Biological Psychiatry: Cognitive Neuroscience and Neuroimaging, 6(6), 607–615. 10.1016/j.bpsc.2020.07.016

Mattingly, D. T., Richardson, M. K., & Hart, J. L. (2024). Prevalence of and trends in current cannabis use among US youth and adults, 2013–2022. Drug and Alcohol Dependence Reports, 12, 100253. 10.1016/j.dadr.2024.100253

McGraw, L. A., & Young, L. J. (2010). The prairie vole: An emerging model organism for understanding the social brain. Trends in Neurosciences, 33(2), 103–109. 10.1016/j.tins.2009.11.006

Mechoulam, R., & Parker, L. A. (2013). The Endocannabinoid System and the Brain. Annual Review of Psychology, 64(Volume 64, 2013), 21–47. 10.1146/annurev-psych-113011-143739

Nashed, M. G., Hardy, D. B., & Laviolette, S. R. (2021). Prenatal Cannabinoid Exposure: Emerging Evidence of Physiological and Neuropsychiatric Abnormalities. Frontiers in Psychiatry, 11. 10.3389/fpsyt.2020.624275

Nguyen, J. D., Aarde, S. M., Vandewater, S. A., Grant, Y., Stouffer, D. G., Parsons, L. H., Cole, M., & Taffe, M. A. (2016). Inhaled delivery of Δ9-tetrahydrocannabinol (THC) to rats by e-cigarette vapor technology. Neuropharmacology, 109, 112–120. 10.1016/j.neuropharm.2016.05.021

Nguyen, J. D., Creehan, K. M., Kerr, T. M., & Taffe, M. A. (2020). Lasting effects of repeated Δ9 -tetrahydrocannabinol vapour inhalation during adolescence in male and female rats. British Journal of Pharmacology, 177(1), 188–203. 10.1111/bph.14856

Nguyen, J. D., Grant, Y., Kerr, T. M., Gutierrez, A., Cole, M., & Taffe, M. A. (2018). Tolerance to hypothermic and antinoceptive effects of Δ9-tetrahydrocannabinol (THC) vapor inhalation in rats. Pharmacology Biochemistry and Behavior, 172, 33–38. 10.1016/j.pbb.2018.07.007

O’Connell, A., Rogers, R., Healy, E. C., Shaikhyousef, M., Regan, C., Douglass, A., Doolan, A., Fitzpatrick, P., Cox, D. W., & Downes, M. (2025). Vaping and Smoking in Pregnancy-A Meta-Analysis of Neurobehavioural Development in Prenatally Exposed Offspring. Acta Paediatrica (Oslo, *Norway*: 1992), 114(12), 3108–3121. 10.1111/apa.70285

Paul, S. E., Hatoum, A. S., Fine, J. D., Johnson, E. C., Hansen, I., Karcher, N. R., Moreau, A. L., Bondy, E., Qu, Y., Carter, E. B., Rogers, C. E., Agrawal, A., Barch, D. M., & Bogdan, R. (2021). Associations Between Prenatal Cannabis Exposure and Childhood Outcomes: Results From the ABCD Study. JAMA Psychiatry, 78(1), 64–76. 10.1001/jamapsychiatry.2020.2902

Pellis, S. M., & Pellis, V. C. (2007). Rough-and-Tumble Play and the Development of the Social Brain. Current Directions in Psychological Science, 16(2), 95–98. 10.1111/j.1467-8721.2007.00483.x

Pertwee, R. G. (2008). The diverse CB1 and CB2 receptor pharmacology of three plant cannabinoids: Δ9-tetrahydrocannabinol, cannabidiol and Δ9-tetrahydrocannabivarin. British Journal of Pharmacology, 153(2), 199–215. 10.1038/sj.bjp.0707442

Preteroti, M., Wilson, E. T., Eidelman, D. H., & Baglole, C. J. (2023). Modulation of pulmonary immune function by inhaled cannabis products and consequences for lung disease. Respiratory Research, 24(1), 95. 10.1186/s12931-023-02399-1

Rice, D., & Barone, S. (2000). Critical periods of vulnerability for the developing nervous system: Evidence from humans and animal models. Environmental Health Perspectives, 108(suppl 3), 511–533. 10.1289/ehp.00108s3511

Roeder, N. M., Penman, S. L., Richardson, B. J., Wang, J., Freeman-Striegel, L., Khan, A., Pareek, O., Weiss, M., Mohr, P., Eiden, R. D., Chakraborty, S., & Thanos, P. K. (2024). Vaporized Δ9-THC in utero results in reduced birthweight, increased locomotion, and altered wake-cycle activity dependent on dose, sex, and diet in the offspring. Life Sciences, 340, 122447. 10.1016/j.lfs.2024.122447

Ross, J. A., & Levy, S. (2023). The Impact of Cannabis Use on Adolescent Neurodevelopment and Clinical Outcomes Amidst Changing State Policies. Clinical Therapeutics, 45(6), 535–540. 10.1016/j.clinthera.2023.03.009

Russo, E. B. (2007). History of Cannabis and Its Preparations in Saga, Science, and Sobriquet. Chemistry & Biodiversity, 4(8), 1614–1648. 10.1002/cbdv.200790144

Schneider, M. (2009). Cannabis use in pregnancy and early life and its consequences: Animal models. European Archives of Psychiatry and Clinical Neuroscience, 259(7), 383–393. 10.1007/s00406-009-0026-0

Schwope, D. M., Karschner, E. L., Gorelick, D. A., & Huestis, M. A. (2011). Identification of Recent Cannabis Use: Whole-Blood and Plasma Free and Glucuronidated Cannabinoid Pharmacokinetics following Controlled Smoked Cannabis Administration. Clinical Chemistry, 57(10), 1406–1414. 10.1373/clinchem.2011.171777

Semple, B. D., Blomgren, K., Gimlin, K., Ferriero, D. M., & Noble-Haeusslein, L. J. (2013). Brain development in rodents and humans: Identifying benchmarks of maturation and vulnerability to injury across species. Progress in Neurobiology, 0, 1–16. 10.1016/j.pneurobio.2013.04.001

Shorey-Kendrick, L. E., Roberts, V. H. J., D’Mello, R. J., Sullivan, E. L., Murphy, S. K., Mccarty, O. J. T., Schust, D. J., Hedges, J. C., Mitchell, A. J., Terrobias, J. J. D., Easley, C. A., Spindel, E. R., & Lo, J. O. (2023). Prenatal delta-9-tetrahydrocannabinol exposure is associated with changes in rhesus macaque DNA methylation enriched for autism genes. Clinical Epigenetics, 15(1), 104. 10.1186/s13148-023-01519-4

Sorkhou, M., Singla, D. R., Castle, D. J., & George, T. P. (2024). Birth, cognitive and behavioral effects of intrauterine cannabis exposure in infants and children: A systematic review and meta-analysis. *Addiction (Abingdon*, England), 119(3), 411–437. 10.1111/add.16370

Spindle, T. R., Cone, E. J., Schlienz, N. J., Mitchell, J. M., Bigelow, G. E., Flegel, R., Hayes, E., & Vandrey, R. (2018). Acute Effects of Smoked and Vaporized Cannabis in Healthy Adults Who Infrequently Use Cannabis: A Crossover Trial. JAMA Network Open, 1(7), e184841. 10.1001/jamanetworkopen.2018.4841

Tadesse, A. W., Ayano, G., Dachew, B. A., Betts, K., & Alati, R. (2025). Maternal perinatal cannabis use disorder and the risk of anxiety disorders in offspring: Insights from a longitudinal data-linkage cohort study. Journal of Affective Disorders, 389, 119743. 10.1016/j.jad.2025.119743

Taffe, M. A., Creehan, K. M., Vandewater, S. A., Kerr, T. M., & Cole, M. (2021). Effects of Δ^9^-tetrahydrocannabinol (THC) vapor inhalation in Sprague-Dawley and Wistar rats. Experimental and Clinical Psychopharmacology, 29(1), 1.

Volkow, N. D., Han, B., Compton, W. M., & McCance-Katz, E. F. (2019). Self-reported Medical and Nonmedical Cannabis Use Among Pregnant Women in the United States. JAMA, 322(2), 167–169. 10.1001/jama.2019.7982

Weimar, H. V., Wright, H. R., Warrick, C. R., Brown, A. M., Lugo, J. M., Freels, T. G., & McLaughlin, R. J. (2020). Long-term effects of maternal cannabis vapor exposure on emotional reactivity, social behavior, and behavioral flexibility in offspring. Neuropharmacology, 179, 108288. 10.1016/j.neuropharm.2020.108288

Young, L. J., & Wang, Z. (2004). The neurobiology of pair bonding. Nature Neuroscience, 7(10), 1048–1054. 10.1038/nn1327

Young-Wolff, K. C., Adams, S. R., Alexeeff, S. E., Zhu, Y., Chojolan, E., Slama, N. E., Does, M. B., Silver, L. D., Ansley, D., Castellanos, C. L., & Avalos, L. A. (2024). Prenatal Cannabis Use and Maternal Pregnancy Outcomes. JAMA Internal Medicine, 184(9), 1083–1093. 10.1001/jamainternmed.2024.3270

Young-Wolff, K. C., Chi, F. W., Campbell, C. I., Alexeeff, S. E., Ansley, D., Vanderziel, A., & Lapham, G. T. (2025). Frequency of Preconception and Prenatal Cannabis Use and Nausea and Vomiting in Pregnancy. Obstetrics and Gynecology, 145(5), 519–522. 10.1097/AOG.0000000000005884

Young-Wolff, K. C., Chi, F. W., Campbell, C. I., Does, M. B., Wysota, C. N., Ansley, D., Castellanos, C., & Lapham, G. T. (2025). Association of preconception cannabis use frequency with cannabis use during early pregnancy. BMC Pregnancy and Childbirth, 25(1), 1044. 10.1186/s12884-025-08190-y

Young-Wolff, K. C., Chi, F. W., Lapham, G. T., Alexeeff, S. E., Does, M. B., Ansley, D., & Campbell, C. I. (2024). Changes in Prenatal Cannabis Use Among Pregnant Individuals From 2012 to 2022. Obstetrics and Gynecology, 144(4), e101–e104. 10.1097/AOG.0000000000005711

Zhou, S., Rosenthal, D. G., Sherman, S., Zelikoff, J., Gordon, T., & Weitzman, M. (2014). Physical, Behavioral, and Cognitive Effects of Prenatal Tobacco and Postnatal Secondhand Smoke Exposure. Current Problems in Pediatric and Adolescent Health Care, 44(8), 219–241. 10.1016/j.cppeds.2014.03.007

Zuardi, A. W. (2006). History of cannabis as a medicine: A review. Brazilian Journal of Psychiatry, 28, 153–157.

